# An antisense oligonucleotide leads to suppressed transcriptional elongation of *Hdac2* and long-term memory enhancement

**DOI:** 10.1101/618116

**Authors:** Shane G. Poplawski, Krassimira A. Garbett, Rebekah L. McMahan, Holly B. Kordasiewicz, Hien Zhao, Andrew J. Kennedy, Slavina B. Goleva, Teresa H. Sanders, S. Timothy Motley, Eric E. Swayze, David J. Ecker, J. David Sweatt, Todd P. Michael, Celeste B. Greer

## Abstract

Repression of the memory suppressor gene <u>h</u>istone <u>d</u>e<u>ac</u>etylase 2 (*Hdac2*) in mice elicits cognitive enhancement, and drugs that block HDAC2 catalytic activity are being investigated for treating disorders affecting memory. Currently available compounds that target HDAC2 are not specific to the HDAC2 isoform, and have short half-lives. Antisense oligonucleotides (ASOs) are a class of drugs that base pair with RNA targets and exhibit extremely long-lasting, specific inhibition relative to small molecule drugs. We utilized an ASO to reduce *Hdac2* messenger RNA (mRNA) quantities, and explored its longevity, specificity, and mechanism of repression. A single injection of the *Hdac2*-targeted ASO in the central nervous system diminished *Hdac2* mRNA levels for at least 4 months in the brain, and knockdown of this factor resulted in significant memory enhancement for 8 weeks in mice. RNA-seq analysis of brain tissues revealed that the ASO repression was specific to the *Hdac2* isoform relative to other classical *Hdac* genes, and caused alterations in levels of other memory-associated mRNAs. In cultured neurons, we observed that the *Hdac2*-targeted ASO suppressed *Hdac2* mRNA and an *Hdac2* non-coding regulatory extra-coding RNA (ecRNA). The ASO not only triggered a reduction in mRNA levels, but also elicited direct transcriptional suppression of the *Hdac2* gene through blocking RNA polymerase II elongation. These findings suggest transcriptional suppression of the target gene as a novel mechanism of action of ASOs, and opens up the possibility of using ASOs to achieve lasting gene silencing in the brain without altering the nucleotide sequence of a gene.

## INTRODUCTION

Long-term memory formation and retention requires coordinated transcriptional changes that are regulated by modifications to the epigenome. Decreasing acetylation by inhibiting histone acetyltransferases (HATs) such as CREB-binding protein (CBP) impairs long-term memory (Alarcon et al. 2004; Korzus et al. 2004; Wood et al. 2005), while increasing acetylation by inhibiting histone deacetylases (HDACs) enhances long-term memory (Levenson et al. 2004; Hawk et al. 2011). Eleven isoforms of classical HDAC proteins exist in mammals, and evidence suggests HDAC2 in particular is responsible for regulating synaptic plasticity and memory formation relative to its close homolog HDAC1 and other HDAC isoforms (Guan et al. 2009). Knockout of the *Hdac2* gene in mice improves hippocampal and prefrontal-cortex dependent learning tasks (Guan et al. 2009; Morris et al. 2013). Specific inhibition of HDAC2 has been a goal of pharmacological design (Wang et al. 2005; Choubey and Jeyakanthan 2018), but a completely selective inhibitor of HDAC2 catalytic activity has remained elusive because of poor pharmacokinetics and promiscuous subtype selectivity.

Antisense oligonucleotides (ASOs) are clinically useful for treating a variety of diseases (Stein and Castanotto 2017). They employ base pairing with a target messenger RNA (mRNA) to achieve selectivity. Therefore, we previously designed and tested an ASO targeting *Hdac2* mRNA. This *Hdac2* ASO elicited substantial memory enhancement in wild-type mice in object location memory tests, and it rescued impaired memory in a mouse model of autism (Kennedy et al. 2016). ASOs targeting other mRNAs reduce expression of their target genes in the central nervous system (Southwell et al. 2014) for months after the last delivery of the drug (Kordasiewicz et al. 2012; Meng et al. 2015a). Because of potential therapeutic utility of the *Hdac2* ASO, we wanted to determine the longevity of its *Hdac2* mRNA reduction and the duration of the elicited cognitive enhancement.

The two most commonly utilized mechanisms for ASOs in therapeutic applications are 1) recruitment of RNaseH1 to the RNA/ASO hybrid and subsequent degradation of the RNA (Wu et al. 2004a), and 2) correction of splicing defects that lead to disease when the ASO is designed to target splice junctions (Sazani et al. 2002; Alter et al. 2006). Because ASOs can modulate splicing, which occurs cotranscriptionally during the synthesis of RNA (Beyer et al. 1981; Osheim et al. 1985; Wu et al. 1991; Merkhofer et al. 2014), we reasoned ASOs could possibly interfere with transcription itself. Therefore, we examined more closely whether ASOs only mediate RNA degradation or might also be involved in preventing their transcriptional synthesis.

We report here that our *Hdac2*-targeting ASO is long-lasting and specific. A single injection of *Hdac2*-targeted ASO in vivo reduced *Hdac2* mRNA for at least 4 months, and increased memory for 8 weeks. It has high selectivity for *Hdac2*, but not other related histone deacetylase isoforms. Furthermore, it affects the expression levels of hundreds of genes in the brain. These genes are involved in extracellular signal-regulated kinase (ERK) signaling, memory-associated “immune” functions, synapse formation, and the regulation of flavonoids. Although the *Hdac2* ASO used herein was designed to mediate degradation of target mRNA, we also found that the ASO elicits repression of an *Hdac2* regulatory RNA, and stimulates transcriptional suppression of its target gene by blocking RNA polymerase II (RNA Pol II) elongation.

## RESULTS

### Long-term target knockdown and behavioral memory enhancement in vivo

Phosphorothiate and 2’-MOE modified ASOs have been shown to elicit prolonged RNA knockdown in the brain (Wu et al. 2004b; Kordasiewicz et al. 2012; Meng et al. 2015b). We wanted to test the longevity of an *Hdac2*-targeting ASO previously shown to achieve potent knockdown (Kennedy et al. 2016), we are calling here *Hdac2* ASO1. A structurally similar scrambled (SCR) ASO that targets no known mouse genes was used as a control. We did a single intracerebroventricular (ICV) injection of SCR ASO or *Hdac2* ASO1 in mice, and examined molecular and behavioral changes out to 40 weeks post-surgery (Fig. 1A). The targeting ASO significantly reduces *Hdac2* mRNA in the cortex for 32 weeks after a single *Hdac2* ASO1 injection (Fig. 1B). HDAC2 protein was significantly lower in the *Hdac2* ASO1 group compared to the scrambled group in cortex from 3 days to 16 weeks after treatment (Fig. 1C). The injection also elicits repression of *Hdac2* mRNA level (Fig. 1D) and HDAC2 protein expression (Fig. 1E) in the hippocampus at 2, 8, and 16 weeks after injection.

**Figure 1.**
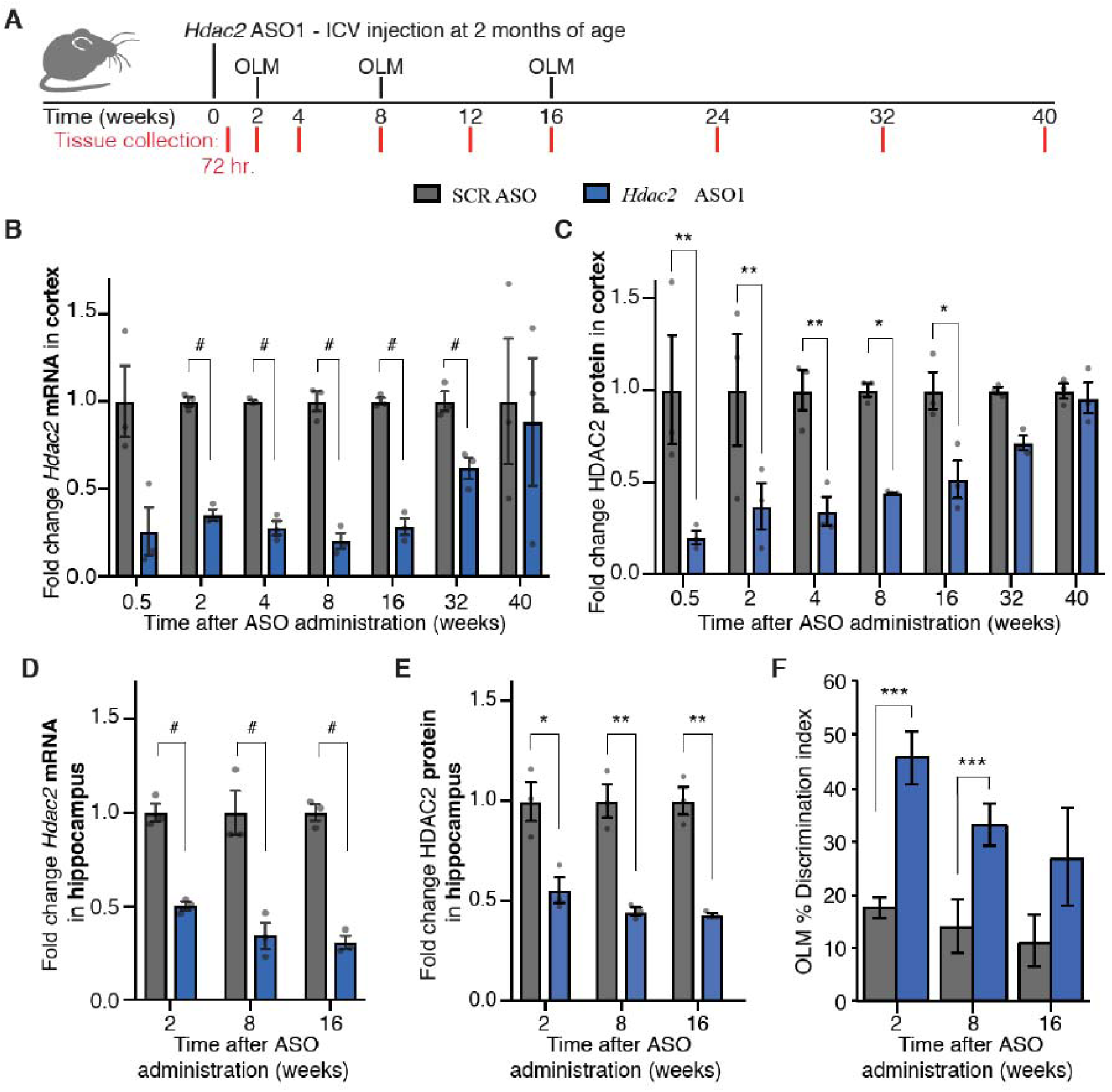
*Hdac2* mRNA, HDAC2 protein. and cognitive enhancement levels across time after ICV injection of ASOs. *(A)* Timeline ofl ongitudinal study. *N* = 3 mice for tissue analysis at each time point. *(B)* Fold change in *Hdac2* mRNA level for *Hdac2* ASOl relative to SCR ASO group in RNA-seq of cortex. Relative expression change in *Hdac2* ASOl group to SCR ASO was calculated from FPKM. (C) Quantitation ofHDAC2 protein in *Hdac2* ASOl group relative to SCR from western blots of co11ical samp les. *(D)* Fold change of *Hdac2* RNA in *Hdac2* ASOl group relative to SCR in RNA-seq of hippocampal samp les. *(E)* Quantitation of ammmt of HDAC2 protein in *Hdac2* ASO1 group relative to SCR from western blots in hippocampal samples. *(F)* Disc1imination index of OLM test for *Hdac2* ASOl and SCR ASO. *N=* 13 12 10, 12 7 and 12 (for each group left to right). Enor bars represent ± SEM. Two-way ANOVA with Sidak’s multiple comparisons *post hoc* test was conducted for western data. Mann-Whitney U test was done for OLM. # = FDR < 0.001. * = *p* < 0.05, ** = *p* < 0.01, *** = *p* < 0.001. Grey dots show values for individual replicates. Enor barn represent ± standard e1rnr of the mean (SEM).

The liver, which is one of the primary peripheral sites of ASO accumulation (Geary et al. 2015), showed no significant mRNA or protein reduction at the two time points tested (2 and 32 weeks) after ICV injection (Supplemental Fig. S1A). Perhaps because of its anatomical distance from the cerebral ventricles, the cerebellum RNA-seq analysis revealed milder, yet significant, knockdown. In the cerebellum, the mRNA knockdown was above 50% and it was statistically significant between 2 and 16 weeks after ICV, However, protein analysis showed no significant downregulation of protein levels at 2 and 32 weeks (Supplemental Fig. S1B).

Blocking HDAC2 has been shown to enhance memory formation, so we tested the duration of spatial memory enhancement in the ICV-injected mice. We used an object location memory (OLM) assessment of the treated animals at weeks 2, 8 and 16. Briefly, the OLM assessment is based on the spontaneous tendency of rodents to spend more time exploring an object that has been relocated. A higher discrimination index indicates that the mouse remembers the familiar placement. Animals that received a single ICV injection of *Hdac2* ASO1 had a higher discrimination index compared to mice that received the control SCR ASO, with statistically significant effects at weeks 2 and 8 and a trend at 16 weeks (Fig. 1F).

### *Hdac2* ASO1 does not cross the brood-brain barrier

Previous reports show that ASOs do not cross the blood-brain barrier (Cossum et al. 1993; Smith et al. 2006), and we wanted to test this for *Hdac2* ASO1. Consistent with prior studies, injection of *Hdac2* ASO1 into the tail vein did not repress *Hdac2* mRNA in the brain. We do see knockdown in the liver at 2 weeks in these intravenously injected animals (Supplemental Fig. S1C).

### Isoform specificity of *Hdac2*-targeted ASOs

Using the RNA-seq data, we examined the isoform specificity of the *Hdac2* ASO in vivo across the longitudinal study. Of the 11 isoforms of classical HDACs, the *Hdac2* gene was the only isoform significantly changed at any time during the 40 weeks of the study in the cortex, hippocampus (Fig. 2A), and cerebellum (Supplemental Fig. S1D).

**Figure 2.**
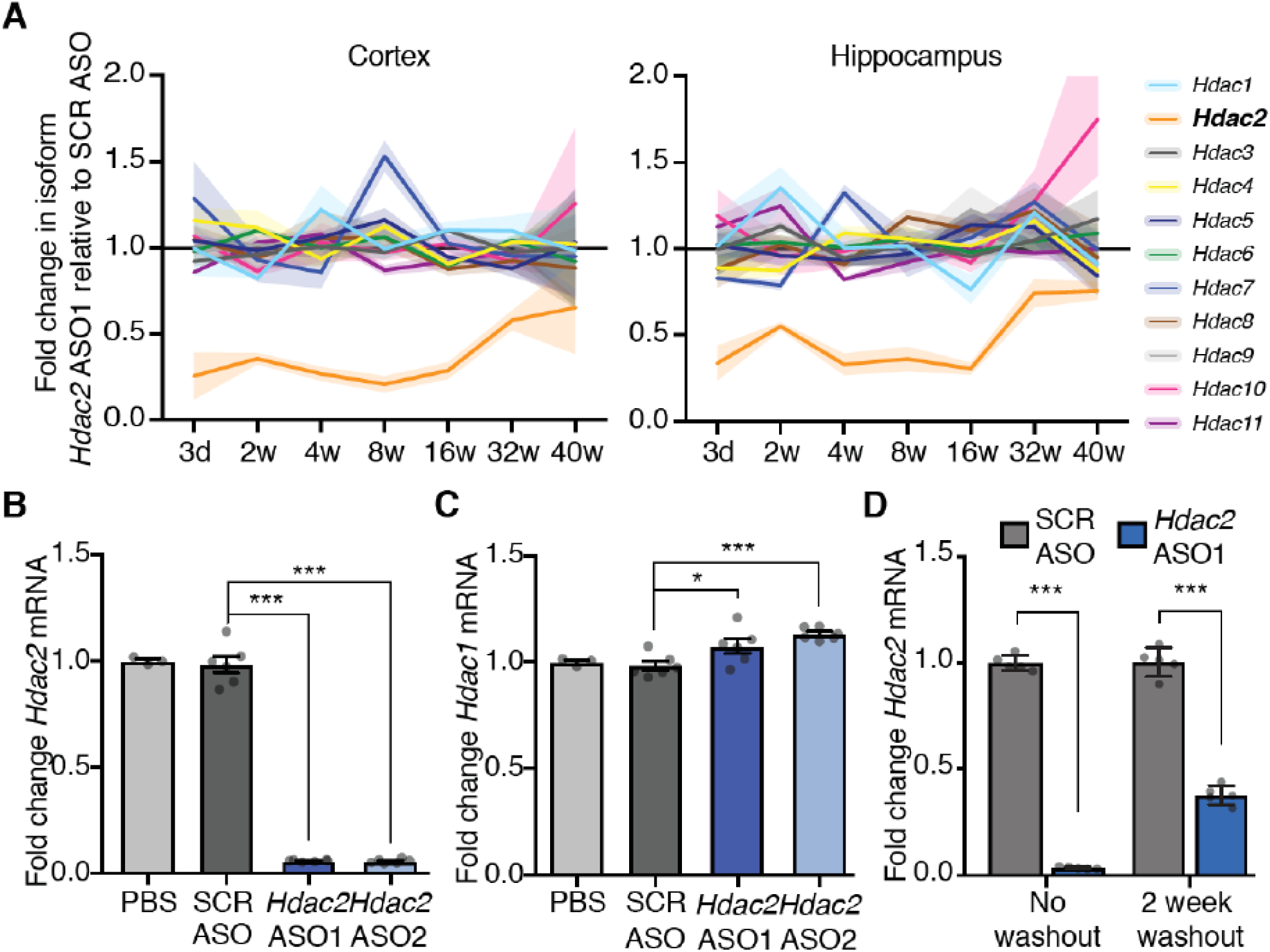
Specificity of Hda c2-target ing ASOs and knockdown efficiency in neurons. *(A)* Isofonn specificity in mouse brain subregions. Fold change calculated from *Hdac2* ASOl FPKM n01malized to average SCR ASO FPKM. Shaded areas represent± SEM. *(B)* Fold change in *Hdac2* mRNA expression after 1-week, 10 pM ASO treatment n01malized to *Hprt* in mouse prima1y cortical neurons. (C) Same samples as used in *B* were tested with *Hdacl* specific primers. *(D)* Comparison of expression of *Hdac2* R A after continuous 16-day treatment or 2-day treatment with ASO cells were rinsed with media, media was replaced and cells were incubated without ASO for two weeks. Enor bars represent± standard enor of the mean (SEM). Grey dots show relative expression values for individual replicates. *N* = 6 wells of cell culture from two biological replicates, each performed with 3 technical replicates. One-way A OVA with Duilllett’ s multiple comparisons *post hoc* test for Band *C* and two-way ANOVA with Sidak’ s multiple comparison s *post hoc* test for *D* were used. * = *p* < 0.05, *** = *p* < 0.001.

Brain tissue contains a variety of cell types, and since the cognitive enhancement functions of *Hdac2* have been ascribed predominantly to gene regulation in neurons (Guan et al. 2009; Penney and Tsai 2014), we next tested for specific and long-lasting ASO-directed *Hdac2* knockdown in neurons. We further wanted to see if our findings are unique to ASO1 or generalizable to another ASO targeting *Hdac2* mRNA at a different site, so we applied *Hdac2* ASO2 alongside ASO1 in mouse primary neuronal culture. ASO1 targets the 3’ untranslated region (UTR), and ASO2 targets exon 10 of the *Hdac2* mRNA. Both *Hdac2* ASOs lead to *Hdac2* mRNA knockdown in neurons after a week of treatment relative to SCR ASO control measured by reverse transcription followed by quantitative PCR (RT-qPCR, Fig. 2B). Furthermore, the two *Hdac2* ASOs show no suppression of closely related *Hdac1* mRNA (Fig. 2C), validating their isoform specificity in neurons. Actually, *Hdac1* was mildly increased in expression, which may be indicative of a compensatory mechanism (El-Brolosy and Stainier 2017). Additionally, we confirmed the efficacy and specificity of the *Hdac2* ASOs in a mouse neuroblastoma Neuro2a (N2a) cell line differentiated with serum-starvation conditions (dN2a, Supplemental Fig. S2A). The *Hdac2* ASOs likewise reduce *Hdac2* (Supplemental Fig. S2B), but not *Hdac1* mRNA in this culture system (Supplemental Fig. S2C). To test if repression of *Hdac2* mRNA by *Hdac2* ASO1 is long-lasting in primary neurons like was observed in the brain in vivo, we tested *Hdac2* mRNA expression 14 days after rinsing and washing out the ASOs, and saw significant reduction persisting after the washout (Fig. 2D).

### Global transcription changes induced by *Hdac2* ASO1 in vivo

Because HDAC2 is a regulator of histone acetylation, chronic reduction of HDAC2 protein is expected to cause changes to the transcriptome. Looking at the ~800 significantly changed genes in RNA-seq between 2 to 8 weeks, when animals had significantly enhanced memory in OLM (Fig. 1F), in the cortex and hippocampus revealed subgroups of genes with expression changes. In both regions, *Hdac2* ASO1 significantly altered the expression of a small set of genes with immune functions and genes with involvement in ERK signaling. Many of these “immune” genes also have known roles in regulating memory (38). Other subgroups of genes that are identified in the cortex (Fig. 3A, blue cluster and red cluster, Table S2) are involved in cell adhesion, transcription, neuron projections, synapse organization, and myelination. The red cluster in Fig. 3B, represents a set of genes altered in the hippocampus involved in glucuronidation and flavonoid biosynthesis (Table S3).

### ASO regulation of a putative *Hdac2* extra-coding RNA

Previous work has demonstrated the existence of extra-coding RNAs (ecRNA) generated from many neuronal protein-coding genes, which promote their transcription (Di Ruscio et al. 2013; Savell et al. 2016). These RNAs are sense transcripts that are unspliced and transcribe over mRNA sequences and prevent repression of their gene of origin. ecRNAs begin transcription upstream of the transcription start site (TSS), and terminatedownstream of the transcription end site (TES) of the gene they regulate. Since ecRNAs contain all of the sequences within the mRNA they control, our *Hdac2* mRNA-targeting ASOs have target sequences matching both mRNA and ecRNA. We know from previous work that *Fos* has an ecRNA regulating its expression, and targeting the ecRNA of *Fos* with an ASO designed against a region downstream of the TES of *Fos* can reduce *Fos* mRNA level (Savell et al. 2016). We likewise found that *Fos* ecRNA ASOs reduce *Fos* mRNA transcript levels (Supplemental Fig. S3A). Moreover, we found that a *Fos* mRNA-targeting ASO downregulates the expression of *Fos* ecRNA as well (Supplemental Fig. S3B). This led us to investigate whether our *Hdac2* mRNA-targeting ASOs also inhibit an *Hdac2* ecRNA.

We first looked for evidence in previously published RNA-seq datasets that the ecRNA is generated from the *Hdac2* gene in primary neuronal cultures of mice and rats. We used an RNA-seq library that detected lowly expressed and transient RNAs (Kim et al. 2010), and a dataset that identified ecRNAs in rat primary neurons. The second dataset has polyadenylated transcripts separated from non-polyadenylated transcripts to better identify the non-polyadenylated ecRNAs (Savell et al. 2016). Reads in regions upstream of the TSS, in introns, and extending beyond the 3’UTR of *Hdac2*, consistent with the presence of an *Hdac2* ecRNA, are seen in primary neurons of mice and rats (Fig. 4A). Furthermore, these *Hdac2* ecRNA reads are predominantly present in the non-polyadenylated fraction (Fig. 4A, PolyA-tracks). These findings suggest an *Hdac2* ecRNA is generated. Because ASO1 and ASO2 target sequence is present in the mRNA, pre-mRNA, and ecRNA (Fig. 4B), we wanted to test if these ASOs repress the ecRNA as well as the mRNA. We found that in primary cortical neurons, ecRNA expression was reduced by both *Hdac2* ASO1 and ASO2 (Fig. 4C), demonstrating that *Hdac2* mRNA-targeting ASOs are also capable of down-regulating the expression levels of the *Hdac2* ecRNA. Likewise, targeting the ecRNA with the ecRNA-specific ASO reduced significantly not only the ecRNA (Fig. 4D, right), but the *Hdac2* mRNA level as well (Fig. 4D, left). From these data, we conclude that the two transcribed products of the *Hdac2* gene, the mRNA and the ecRNA, are both efficiently downregulated by ASOs, and solely targeting the ecRNA is sufficient to elicit mRNA knockdown.

**Figure 4.**
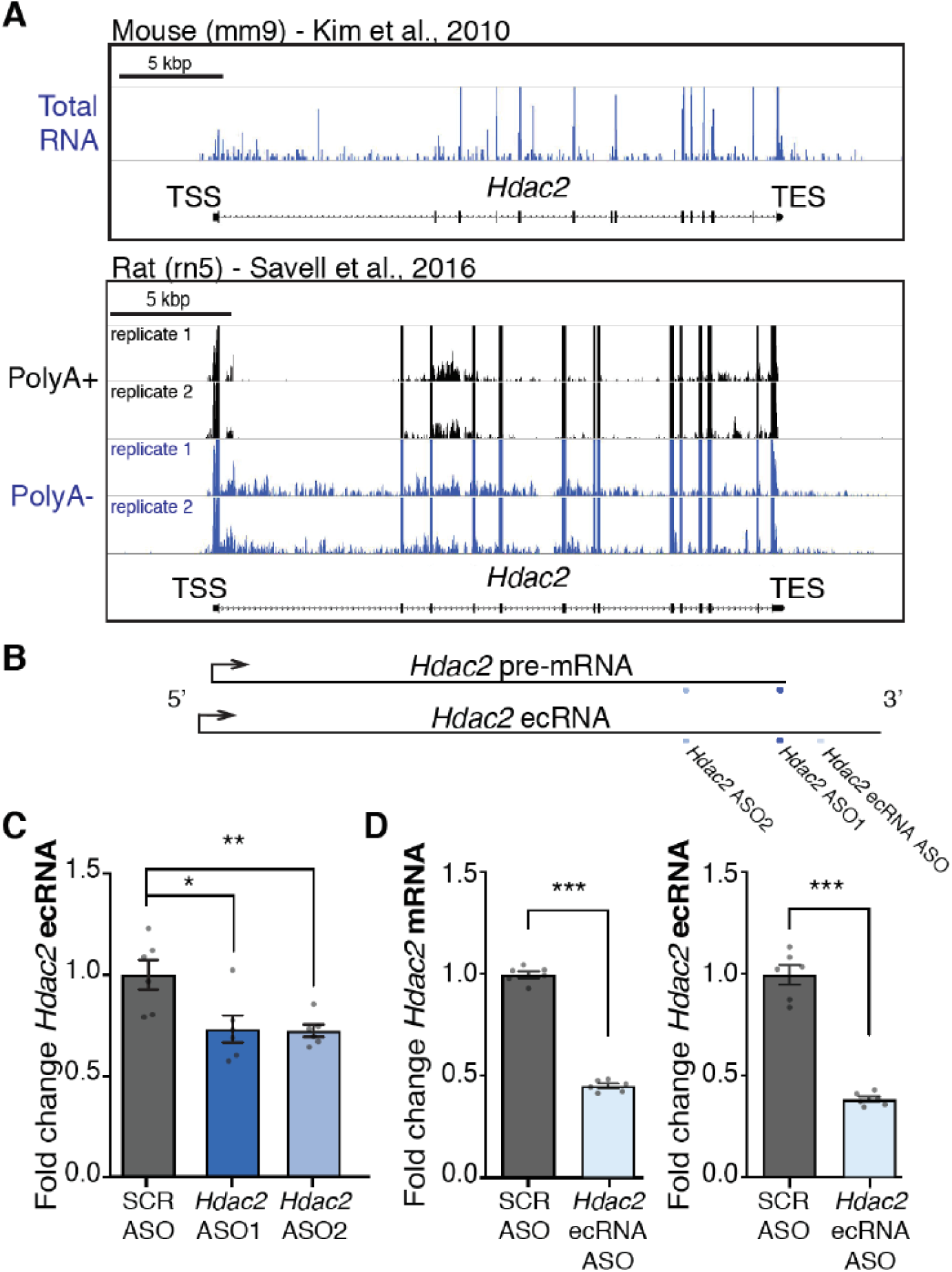
*Hdac2* has a putative ecRNA and it is repressed by *Hdac2* ASOs. *(A)* RNA-seq reads at *Hdac2* gene from indicated sequencing libraries. Reads were stranded, and only the reads representing sense transcripts relative to the direction of the *Hdac2* gene are shown (+ for mouse, - for rat). Gene tracks for rat were flipped to show the 5’ end of *Hdac2* on the left. *(B)* Diagram of *Hdac2* pre-mRNA and ecRNA, with location of targeting sequences for *Hdac2* ASOs. (C) Signal from RNA generated beyond the TES was detected with RT-qPCR using primers designed to the 3’ end of the *Hdac2* ecRNA (TES +1.4 kilo-base pairs (kbp)). Reverse transcription of RNA was done with random primers. *N* = 6 wells of cell culture from 2 biological replicates done in triplicate. *(D)* The *Hdac2* ecRNA ASO repressed *Hdac2* mRNA (left) and *Hdac2* ecRNA (right, ecRNA primer targets TES +1.4 kbp). *N* = 6 wells of cell culture from 2 biological replicates done in triplicate. *Hdac2* mRNA and ecRNA signal was 1101malized to *Hprt.* En-or bars represent ± SEM. Grey dots show values for individual replicates. * _=_ *p* < 0.05, ** _=_ *p* < 0.01, *** _=_ *p* < 0.001 by one-way ANOVA with Dunnet’s multiple comparisons *post hoc* test in C, or Students’ *t* tests in *D.*

### *Hdac2* ASO1 causes no detectable change to DNA methylation at *Hdac2* gene

ecRNAs are reported to interact with DNA methyltransferases (DNMT), and prevent DNA methylation to maintain transcriptional accessibility (Di Ruscio et al. 2013; Savell et al. 2016). These prior studies show DNMT1 and DNMT3a are suppressed by ecRNA binding. Contrary to these models, we find that DNMT inhibitors 5-azacitidine (5-Aza) and RG108, which target both DNMT1 and DNMT3a isoforms, do not antagonize the repression of *Hdac2* mRNA by *Hdac2* ASO1 (Supplemental Fig. S3C). In addition, RG108 does not overcome the suppression of *Fos* mRNA and *Fos* ecRNA by *Fos* ecRNA ASOs (Supplemental Figs. S3D and S3E), suggesting ASO-mediated repression of *Fos* is not DNA methylation-dependent either. Furthermore, direct measures of DNA methylation across the *Hdac2* gene after treatment with *Hdac2* ASO1 measured by MeDIP-qPCR also do not reveal any DNA methylation changes (Supplemental Fig. S3F).

### Direct transcriptional suppression by ASOs

Since ecRNAs are reported to regulate transcriptional accessibility, we tested the hypothesis that ASOs trigger transcriptional suppression of *Hdac2* mRNA production. To determine if ASO1 and ASO2 prevent *Hdac2* pre-mRNA production, we conducted nuclear run-on (NRO) experiments to quantify nascent *Hdac2* transcripts. During NRO, newly synthesized transcripts are purified by immunoprecipitation (IP). The IP method we optimized was able to isolate BrU-labeled RNA (+ samples), while washing away unlabeled RNA (-samples) with very low background signal (Fig. 5A). The level of *Hdac2* pre-mRNA in the immunoprecipitated nascent RNA fraction was measured by RT-qPCR. We observed that *Hdac2* ASO1 treatment results in a significant reduction of transcription towards the 3’ end of the *Hdac2* gene in the 3’UTR in primary neurons (Fig. 5B). *Hdac2* ASO2, although it targets a different exon, repressed *Hdac2* transcription in a similar pattern to ASO1 in dN2a cells (Fig. 5C). Interestingly, transcription near the 3’, but not the 5’, end of the *Hdac2* gene was impeded in the NRO assay, which suggests that the elongation, but not the initiation, of transcription is blocked by *Hdac2* ASOs. *Hdac2* ASO1 elicited repression across all exons relative to SCR ASO in our sequencing of mature mRNA transcripts (Supplemental Fig. S4A), suggesting this pattern of repression is a unique feature of nascent RNAs. HDACs have been implicated in transcriptional suppression as well as activation (Kim et al. 2013; Greer et al. 2015), so we tested if the changes we observed in the NRO experiment were an indirect effect of blocking HDAC activity using two broad-spectrum HDAC inhibitors, sodium butyrate (NaBu) and Suberoylanilide Hydroxamic Acid (SAHA). There was no repression of *Hdac1* or *Hdac2* transcript levels (Supplemental Fig. S4B), and *Hdac2* transcription in NRO assays was not blocked by NaBu (Supplemental Fig. S4C). Treatment of the cells with SAHA also did not reduce *Hdac2* transcript levels, but actually increased *Hdac1* and *Hdac2* mRNA expression (Supplemental Fig. S4D). Therefore, we conclude that *Hdac2* ASOs directly reduce the transcription of their target gene irrespective of any epigenetic changes resulting from inhibition of deacetylation.

**Figure 5.**
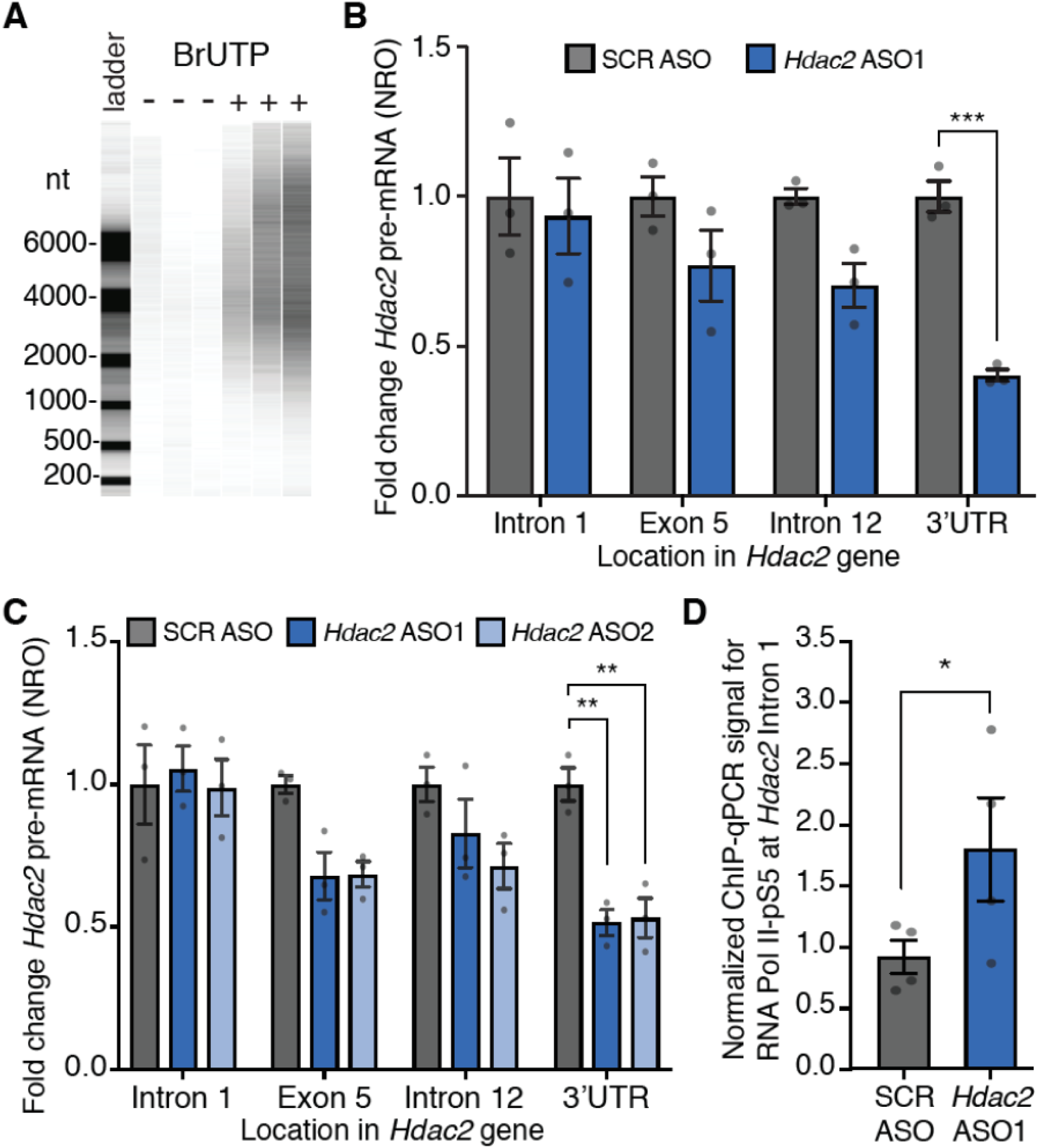
Direct repression of *Hdac2* transcription by *Hdac2* ASOs. *(A)* Bioanalyzer high sensitivity RNA chip gel-like image of BrUTP antibody immunoprecipitated NRO samples made with unlabeled UTP (-) or with BrUTP (+). Units for ladder are shown in nucleotides (nt). *(B)* RT-qPCR of NRO samples made from primary cortical neurons treated with ASOs. *N=* 3 biological replicates from independent preparations of primary neurons. (C) RT-qPCR of NRO samples made from dN2a cells treated with ASOs. *N-* 3 technical replicates. *(D)* ChIP-qPCR with RNA Pol II pS5 antibody was conducted in primary neurons treated with SCR ASO and *Hdac2* ASOl. *N* = 4 biological replicates from independent preparations of primary neurons. Paired Students’ *t* test. Two-way ANOVA with Sidak’s multiple comparisons *post hoc* test was performed for panels *B* and *C*. Err or bars represent ± SEM, grey dots show values for individual replicates.

Based on the NRO assay, we predicted that RNA Pol II might be stalled near the promoter of the *Hdac2* gene, preventing production of full-length transcripts. Therefore, we tested if *Hdac2* mRNA expression is regulated by transcriptional pause release since not all genes are under the control of this mechanism of transcriptional regulation (Greer et al. 2015; Zocchi et al. 2018). Positive transcription elongation factor b (P-TEFb) is an important regulator of transcriptional pausing, and it is inhibited by the small molecule inhibitor Flavopiridol (FLAVO) (Chao and Price 2001). In primary neuronal cultures and dN2a, FLAVO inhibits *Hdac2* mRNA expression (Supplemental Fig. S4E), so we conclude that RNA Pol II pausing is an important regulatory step in *Hdac2* mRNA expression levels. Next, we looked to see if RNA Pol II is stalled at the promoter after *Hdac2* ASO1 treatment. Since in the NRO experiments, the decrease in transcripts occurred downstream of *Hdac2* intron 1, we looked for an accumulation of the initiated form of RNA Pol II (RNA Pol II phosphorylated at serine 5 of the C-terminal domain repeat region; RNA Pol II-pS5) (Komarnitsky et al. 2000; Phatnani and Greenleaf 2006) at intron 1 of the *Hdac2* gene using ChIP-qPCR. Indeed, we found more binding of this active form of RNA Pol II at this pause site in the *Hdac2* gene (Fig. 5D), indicating that *Hdac2* ASO1 leads to a pause in transcription near intron 1 of the *Hdac2* gene.

## DISCUSSION

Our studies indicate that *Hdac2* ASOs provide a powerful avenue to generate long-lasting beneficial changes in epigenomic organization in the central nervous system. Specifically blocking *Hdac2* expression with this high precision compound leads to cognitive enhancement in WT mice, and prior studies suggest specific targeting of *Hdac2* could be beneficial for treatment of autism (Kennedy et al. 2016) and Alzheimer’s disease (Graff et al. 2012).

We identified changes in the expression of many genes implicated in memory formation. For example, we see altered expression of genes associated with ERK signaling in the hippocampus and cortex, which has an important role in learning and memory. This signaling pathway is necessary for long-term potentiation of synaptic activity (Peng et al. 2010). Additionally, we saw alterations in genes first identified as performing immune functions in peripheral tissues, but are also known to promote cognition (Nelson et al. 2013; Ru and Liu 2018) and synaptic plasticity (O’Brien et al. 1999; Golan et al. 2004; Stevens et al. 2007; Fuerst et al. 2008). Examples of immune-related factors with connections to memory we see changed by *Hdac2* ASO1 in our datasets include Down syndrome cell adhesion molecule (DSCAM) proteins, pentraxins, tumor necrosis factor alpha, the complement system, and MHCII. *Hdac2* regulates neuronal spine density, and the HDAC2 protein binds to the promoters of genes involved in synapse formation and plasticity (Guan et al. 2009). Accordingly, we identified that the *Hdac2* ASO affects the expression of genes involved with cell adhesion and synapse formation in the cortex. We also found a subgroup of genes involved in flavonoid regulation in the hippocampus affected by *Hdac2* ASO1 treatment. Importantly, flavonoids are now being studied with regards to their role in cognitive enhancement and regulation of inflammation (Bakoyiannis et al. 2019), suggesting there may be a link between the action of these compounds and *Hdac2*.

*Hdac2* ASOs achieve their knockdown by preventing RNA Pol II elongation (Fig. 6). This transcriptional inactivation may explain the long-term effects of a single application of ASO, and is through an ecRNA-dependent, DNA methylation-independent mechanism. Peptide nucleic acids (PNAs) that base pair with DNA, but have a peptide backbone, have been designed to target gene promoters and prevent RNA Pol II transcription through steric hindrance of transcription factor binding (Wang and Xu 2004). Our data demonstrate that ASOs also act on transcription even if they are designed to target sequences far downstream of the promoter. It will be of interest to discriminate whether ASOs act like PNAs in blocking the binding of transcriptional regulators.

**Figure 6.**
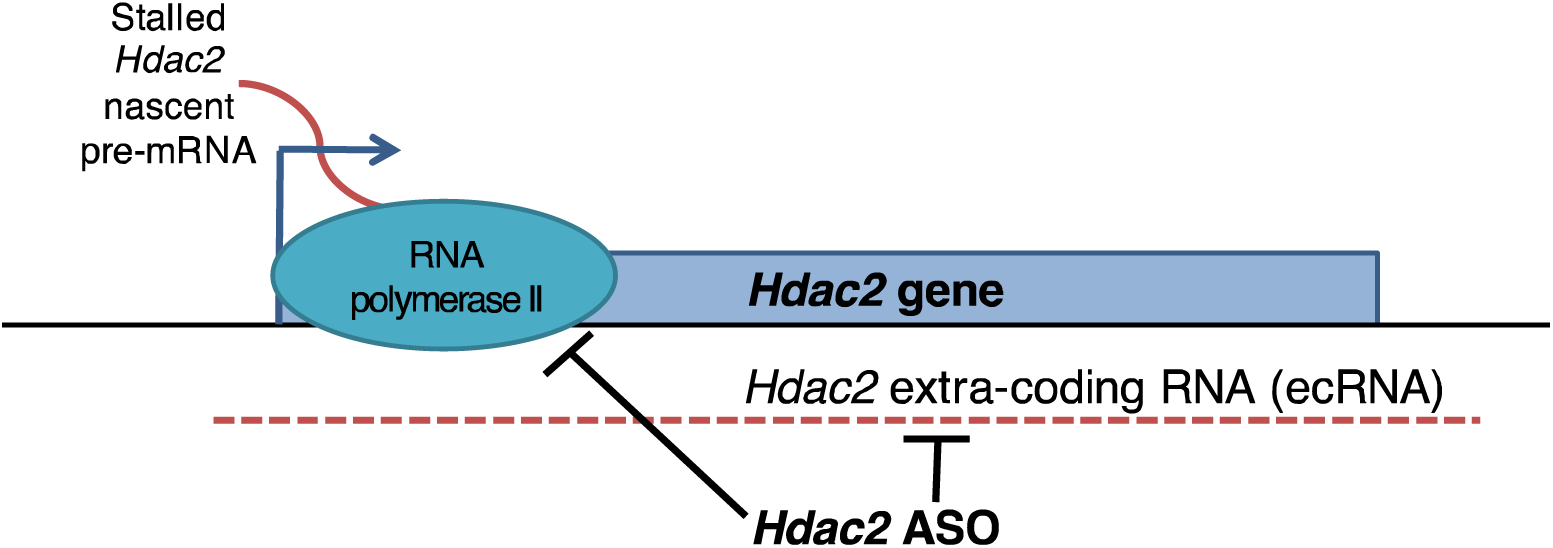
ASOs block transcription by suppressing elongation and repressing ecRNA.

ASOs employ several mechanisms to modulate RNA stability, splicing, and translation (DeVos and Miller 2013). Our study reveals a yet another mechanism of directly blocking transcriptional progression across the gene which enables ASOs modulate mRNA generation. ASOs elicit targeted mRNA reduction that is potent and long lasting without altering the underlying gene sequence, which makes them more attractive than other gene therapy approaches, like CRISPR (Gaj et al. 2013), in certain contexts. This is because the transcriptional changes induced by ASOs are extremely specific, and enduring, but not permanent or damaging to genomic sequences.

## METHODS

### ASOs

*Hdac2* ASO1 (5′-CToCoAoC<u>TTTTCGAGGT</u>ToCoCTA-3′), *Hdac2* ASO2 (5′-AToGoCoA <u>GTTTGAAGTC</u>ToGoGTC-3′), *Hdac2* ecRNA ASO (5′-CCoCoAoA<u>ATCACCTGTT</u>CoToGAA-3′), and non-targeting scrambled (SCR) ASO (5′-GToToToT<u>CAAATACACC</u>ToToCAT-3′) were generated by IONIS using the phosphorothioate and 2′-MOE modified ASO platform. *Fos* ASOs (*Fos* mRNA ASO (5’-UCUGU<u>CAGCTCCCTC</u>CUCCG-3’), *Fos* ecRNA ASO1 (5’-AGAUU<u>GGCTGCTTGG</u>UGGGU-3’), *Fos* ecRNA ASO2 (5’-ACUAG<u>CGTGTCCTCTG</u>AGUGA-3’), and non-targeting SCR ASO (5’-GUUUU<u>CAAATACACC</u>UUCAU-3’)) were ordered from IDT. Sequences are designed for targeting mouse transcripts. Underlined residues are deoxynucleosides, and all others are 2’-*O*-methoxyethyl (2’-MOE) nucleosides. All linkages are phosphorothioate except those indicated by “o” between residues, which are phosphodiester.

### Cell culture

Primary cortical neuron cultures were made from P0 mice. Dissected cortices were treated with papain supplemented with cysteine, and triturated to dissociate neurons. Cells were passed through a 70 μm filter (Falcon), and grown in neurobasal complete media (neurobasal with 1x B27 supplement, 1mM sodium pyruvate, 1mM HEPES, 100U/mL penicillin, 100μg/mL streptomycin, and 0.5mM L-glutamine). Cells were treated overnight with FdU to kill dividing cells on day in vitro (DIV) 3, and 10μM ASO was applied on DIV 5. For washout experiments, neurons were treated 2 days with 10μM ASO. Cells were rinsed with complete neurobasal media, left a few minutes, and media was replaced again. ½ media changes were done every 2 to 3 days for all primary culture experiments. New media for changes did not contain ASO.

Neuro2a (N2a) cells were obtained from ATCC and grown according to recommended conditions. For long treatments plates were coated in poly-L-lysine. Cells attached overnight, and media was changed to differentiation media (DMEM with L-glutamine without glucose, 10mM galactose, 100U/mL penicillin, 100μg/mL streptomycin, and 1x N2 supplement) to make differentiated N2a (dN2a). After 4 days, half the media was changed and supplemented with complete neurobasal media at a ratio of 1:400 neurobasal to differentiation media. Half media changes were done as needed. Replacement media contained drug at the same concentration as the initial treatment.

For *Fos* ecRNA experiments, N2a were transfected with GenMute reagent according the manufacturer’s specifications with ASO at a final concentration of 60nM. Media was changed to differentiation media 5 hours after transfection, and 2 days later changed to neurobasal media. RNA was extracted the following morning.

### Mice

Male B6129S F1 hybrid mice at 2 months of age were acquired from The Jackson Laboratory. All procedures were performed with Institutional Animal Care and Use Committee (IACUC)-approved protocols and conducted in full compliance with the Association for Assessment and Accreditation of Laboratory Animal Care (AAALAC).

### In vivo ASO administration

ASOs were injected by unilateral ICV bolus injection of 300 μg. Mice were anaesthetized with 2% isoflurane and secured in a stereotaxic frame (David Kopf Instruments). ASOs were diluted to 60 μg/μl in saline and injected 15 mg/kg into the lateral ventricle (anterior/posterior [A/P], −0.2; medial/lateral [M/L], −1.0; dorsal/ventral [D/V], −2.4 to the bregma) of 2-month-old mice at a rate of 1 μl/min. After the injection, the needle was kept in place for 5 min., followed by suturing of the incision. Intravenous (IV) injections of ASO (300 μg) were done into the tail vein. 3 animals per group per time point were treated.

### Targeted gene expression analysis

For mouse tissue samples, total RNA and DNA was extracted with AllPrep® DNA/RNA/miRNA kit (Qiagen). RNA was extracted from the right hemisphere of the brain. Total mRNA was reverse transcribed using the iScript cDNA Synthesis Kit (Bio-Rad). For culture samples the RNeasy plus kit (QIAGEN) and SuperScript VILO (Invitrogen) was used according to manufacturer’s instructions. qPCR was performed with the CFX96 Optical Reaction Module (Bio-Rad) using SYBR green (Bio-Rad). Relative gene expression from in vivo samples was determined using the ΔΔCt method (Livak and Schmittgen 2001) and normalized to a housekeeping gene. qPCR primer sequences are listed in Table S1.

### Western blots

Tissue from the left hemisphere of the brain was homogenized in RIPA buffer. Protein samples were run on 4-20% TGX Gels (Bio-Rad) and then transferred to PVDF membranes (Millipore) using standard protocols. Primary antibodies were: HDAC2 (Abcam ab12169) and actin (Abcam ab3280). Secondary antibodies were goat anti mouse IR 680 (LiCor #926-68020) and goat anti mouse IR 800 (LiCor #926-32210). Membranes were imaged on the LiCor Odyssey fluorescence imaging system.

### Object location memory test (OLM)

Mice were habituated to an opaque polyurethane open box (10 × 10 × 12 in. (x, y, z)) containing autoclaved bedding with one black line spatial cue for 3 days (5 min. per day) prior to training. Mice were trained for 10 min. with two 50 ml beakers in a particular location. 24 hr. after training, one beaker was moved to a novel location and the mice were tested for 5 min. Videos were scored by hand and blinded to subject identity. Object interaction was scored as previously described (Haettig et al. 2011).

### Chromatin immunoprecipitation (ChIP)

RNA Pol II ChIP cells were cross-linked with 0.5% formaldehyde in neurobasal for 10 minutes at RT. The crosslinking was stopped with glycine, and cells were immediately placed on ice and lysed in L1 buffer.

Purification of chromatin was done as previously described (Greer et al. 2015). Chromatin was sonicated in Diagenode mini water bath to 100-400 bp fragments. 5μg of antibody 3E8 (Millipore), 50μL of protein G coated dynabeads, and 68μg of chromatin were used per IP. Signal from Intron 1 primers was standardized to input, normalizing to *Gapdh* intron primers. Fold change was calculated relative to SCR ASO signal from *Hdac2* exon 5 primer from each batch of chromatin. ½ of the plates were treated with SCR ASO and *Hdac2* ASO1 for each preparation, so batch effects could be taken into account.

### Methyl-DNA immunoprecipitation (MeDIP)

DNA was isolated with DNeasy blood and tissue kit (QIAGEN) and sonicated to 300-1000bp with a Diagenode bioruptor for 10-16 rounds of 15s on 90s off at a concentration of 20ng/μL in 300μL volume. 3-5μg DNA, 30μL prewashed protein G dynabeads, and 10μg EpiGentek OptimAb 5-Methylcytosine (33D3) monoclonal antibody were used per pull-down. Pull down was done in IP buffer (5mg/mL BSA and 0.05% Triton X-100 in 1x PBS) at 4 °C for 2 hours with rotation. Beads were washed three times with IP buffer on a magnetic stand. DNA was dissociated from beads with 1mg/mL proteinase K overnight at 50°C with shaking, extracted with phenol/chloroform, precipitated in NaCl and ethanol, washed in 70% ethanol, and resuspended in TE buffer.

### Nuclear run-on (NRO)

This procedure is based on several sources (Core et al. 2008; Kim et al. 2013; Greer et al. 2015; Roberts et al. 2015) and optimized to reduce non-specific RNA binding. Nuclei were extracted as previously described (Greer et al. 2015), with lysis buffer containing Igepal concentration optimized based on cell type - 0.25% for dN2a and 0.5% for primary neurons. The run-on reaction was done as previously described (Greer et al. 2015). RNA concentration was measured by nanodrop and normalized. 30μL Protein G dynabeads were washed twice in BrU binding buffer, rotated at RT with 2μg Anti-BrdU antibody (Santa Cruz Biotech, IIB5, sc-32323) in BrU binding buffer for 10 min., blocking buffer was added, and beads rotated another 30 min. at RT. After blocking, beads were washed 2 times with binding buffer. The blocked bead mixture was combined with RNA sample, and put on a rotating stand for 30 min. at RT. After binding, beads were washed twice for 2 min. in BrU binding buffer, once in low salt buffer, once in high salt buffer and twice in TET buffer. Buffer compositions were previously published (Core et al. 2008). On the final TET wash, beads were moved to a new tube, TET was removed, and TRIzol was used to elute and purify RNA as previously described (Roberts et al. 2015). Three rounds of immunoprecipitation were conducted on each sample. Purified RNA samples were heated at 65°C for 5 min. then placed on ice at least for 2 min. prior to IP or reverse transcription reaction. Multiscribe reverse transcriptase was used to make cDNA according to manufacturer’s recommendations.

For the non-specific binding check, NRO samples made with UTP were run in parallel to samples made with BrUTP. The elution after the third round of BrdU immunoprecipitation was run on a bioanalyzer Eukaryote Total RNA Pico Series II chip according the manufacturer’s instructions. Signal intensity is normalized across all samples in the gel-like output image.

### Total RNA-seq

Total RNA-seq libraries were prepared using the TruSeq Stranded Total RNA Library Prep Kit with Ribo-Zero Gold (Illumina) according to manufacturer’s instructions. 1µg of RNA was used as starting material and amplified with 12 PCR cycles. Library size distribution was checked with an Agilent 2100 Bioanalyzer and quantity was determined using qPCR. Libraries were sequenced on an Illumina HiSeq 2500 using a 50-cycle rapid run kit or an Illumina NextSeq instrument using a 75-cycle high throughput kit. Reads were aligned to the GRCm38.p3 mouse genome and transcriptome using TopHat (Trapnell et al. 2009) and differential expression tests were performed using featureCounts (Liao et al. 2014) and edgeR (Robinson et al. 2010; McCarthy et al. 2012) with standard settings. DAVID (Huang da et al. 2009b; Huang da et al. 2009a) was used for functional annotation of genes. Heatmaps were generated with gplots package in R.

### Statistics

ANOVA and Student’s *t* tests were conducted in GraphPad Prism version 8 with indicated post-hoc tests.

## Supporting information

Hippocampus significantly changed genes

Cortex significantly changed genes

## DATA ACCESS

All raw sequencing data generated in this study have been submitted to the NCBI Gene Expression Omnibus (GEO; http://www.ncbi.nlm.nih.gov/geo/) under accession number GSE124726. All relevant data from this study are available on request from the corresponding authors (C.B.G. or T.P.M.).

## ACKNOWLEDGEMENTS

We would like to thank Tim Broderick and Bruce Howard for many helpful discussions over the course of this project. The authors sincerely appreciate help from Jane Wright, Garrett Kaas, and Joseph Weiss with primary neuron generation and isolation, and help from Colin Fricker and Hero Haji with qPCR. We thank Benjamin Coleman and Joseph Weiss for their efforts in preliminary data generation.

## AUTHOR CONTRIBUTIONS

C.B.G., S.G.P., K.A.G., A.J.K., T.P.M., S.T.M. aided with experimental design and analysis. S.G.P. conducted sequencing prep and bioinformatics. C.B.G. conducted in vitro assays and analysis of RNA-seq data. K.A.G. was project manager for Vanderbilt’s team, and conducted the tail vein injection experiments. A.J.K. did ICV infusions. A.J.K and R.L.M. aided with expression analysis, animal behavior, and colony maintenance. R.L.M. conducted immunoblotting. H.B.K. and E.E.S. designed and developed the *Hdac2*-targeting ASOs. S.B.G. redesigned and conducted experiments with the mouse *Fos* ASOs. S.G.P., J.D.S., T.P.M., and C.B.G. wrote the manuscript with assistance from all authors.

