## Supplemental Figures for "An antisense oligonucleotide leads to suppressed transcriptional elongation of *Hdac2* and long-term memory enhancement"

Supplementary Materials:

**Table S1.** qPCR primer sequences

| **Target** | **Forward Primer Sequence (5’ to 3’)** | **Reverse Primer Sequence (5’ to 3’)** |
| --- | --- | --- |
| *Fos* ecRNA | TGCACTTGAGGTCATGGGAC | CTGGCAAGATTGGCTGCTTG |
| *Fos* mRNA | GGGAGCTGACAGATACACTCC | GCAATCTCAGTCTGCAACGC |
| *Gapdh* Intron | CCATAGGTGTGGAGAACCTGC | TGTCTGCATAAAGGCTTGTACCT |
| *Hdac1* mRNA | CACTGTAAGACCACTGCACTA | TGAACTACCCACTGCGAGA |
| *Hdac2* mRNA | AGAAGGAGACAGAGGACAAGA | GAGTCAAATTCAAGGGTTGCTG |
| *Hdac2* Promoter | TAAGACCGAGGGGTGAACCT | GGGTAGTCACACACAGTCCG |
| *Hdac2* Intron 1 | ACCGCTCATGCTTACTGTTT | TCCCCTTAACCCCATCCCAT |
| *Hdac2* Exon 5 | CAATTGGGCTGGAGGACTACAT | AATTCGAGGATGGCAAGCACAA |
| *Hdac2* Intron 12 | AGCATGCCATTACAGCACCT | AGGCTACTGCCCACTCCTTA |
| *Hdac2* 3’UTR | GCCAAGTCAGAACAACTCAGC | ATGTCCTCAAACAGGGAAGGT |
| *Hdac2* TES+0.9 kbp | AAACCTGGTTCTTGCCCTGT | GAATGCGGAATGCCAAAGCA |
| *Hdac2* TES+1.4 kbp | TGTGATCTGCCTTCTGGCAC | ACCAGAACCTCCCAGATCCA |
| *Hdac2* TES+1.7 kbp | CTGTCGGGACAAGGGTTGAA | ATGTGGGGTTCGGTGTCAAG |
| *Hprt* mRNA | GGAGTCCTGTTGATGTTGCCAGTA | GGGACGCAGCAACTGACATTTCTA |
| *Hprt* for NRO | TTGCTCGAGATGTCATGAAGG | TGTAATCCAGCAGGTCAGCAAA |

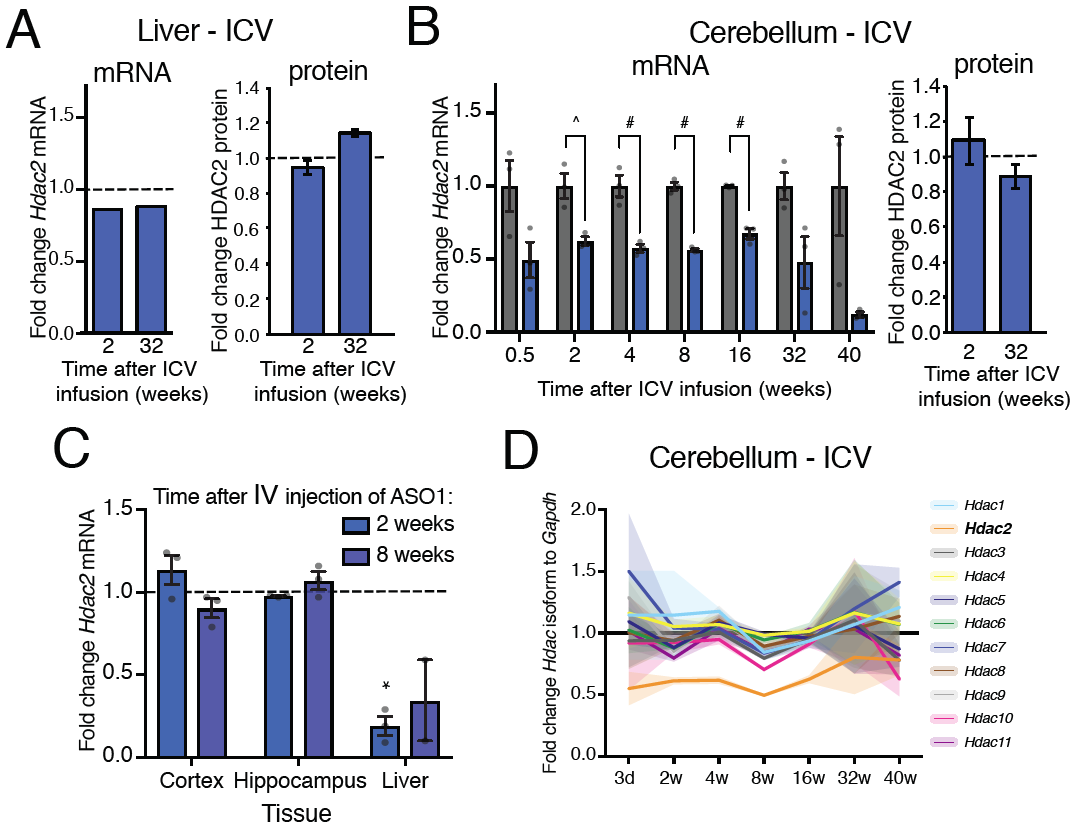

**Fig. S1. Expression of *Hdac2* RNA across the gene and in different tissues with different administration strategies.** (**A**) Fold change of *Hdac2* mRNA measured by RNA-seq and HDAC2 protein level measured by western in the liver. (**B**) Fold change in RNA-seq signal relative to SCR in cerebellum after ICV administration. Western quantitation in cerebellum at 2 and 32 weeks after ASO administration via ICV injection. ^ = FDR < 0.05, # = FDR < 0.001. *N* = 3 animals. (**C**) Expression of *Hdac2* in brain tissues and liver 2 and 8 weeks after IV administration. *N* = 2-3 animals. * = *p* < 0.05 in Student’s *t* test. (**D**) Isoform specificity of ASO1 in cerebellum. Fold change to SCR ASO, normalized to *Gapdh*. Error bars represent ± SEM on RNA graphs, and ± standard deviation on western graphs. Grey dots show values for individual replicates.

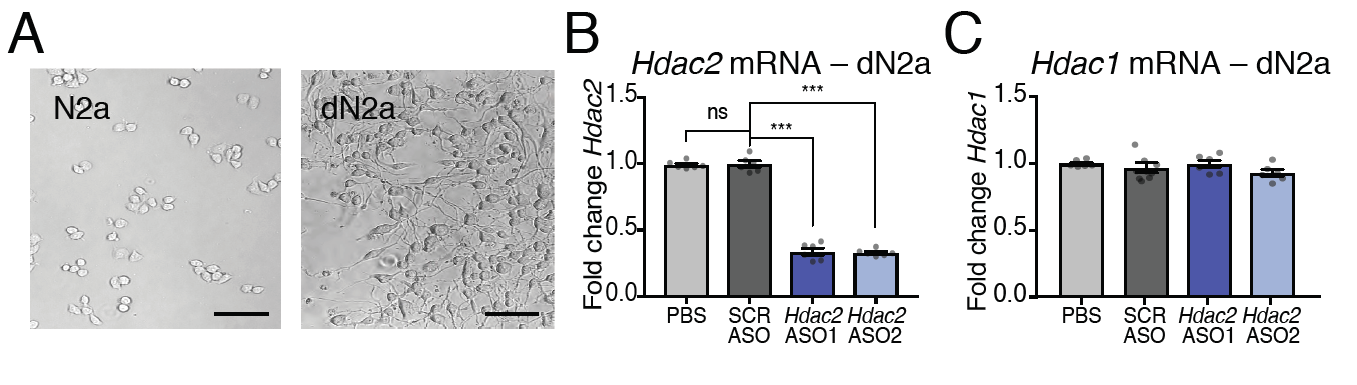

**Fig. S2. ASO specificity in another cell culture system.** (**A**) Bright field images of N2a after plating, then after differentiation into dN2a. Scale bar represents 100μm. (**B**) *Hdac2* expression level from total RNA of dN2a after 1-week treatment with ASO. Relative to *Hprt*. *N* = 6 wells of cell culture from two biological replicates performed in triplicate. (**C**) *Hdac1* expression level in the same samples as panel B. Bars represent ± SEM, grey dots show values for individual replicates. One-way ANOVA, with Dunnett’s multiple comparisons *post hoc* test. *** = *p* < 0.001.

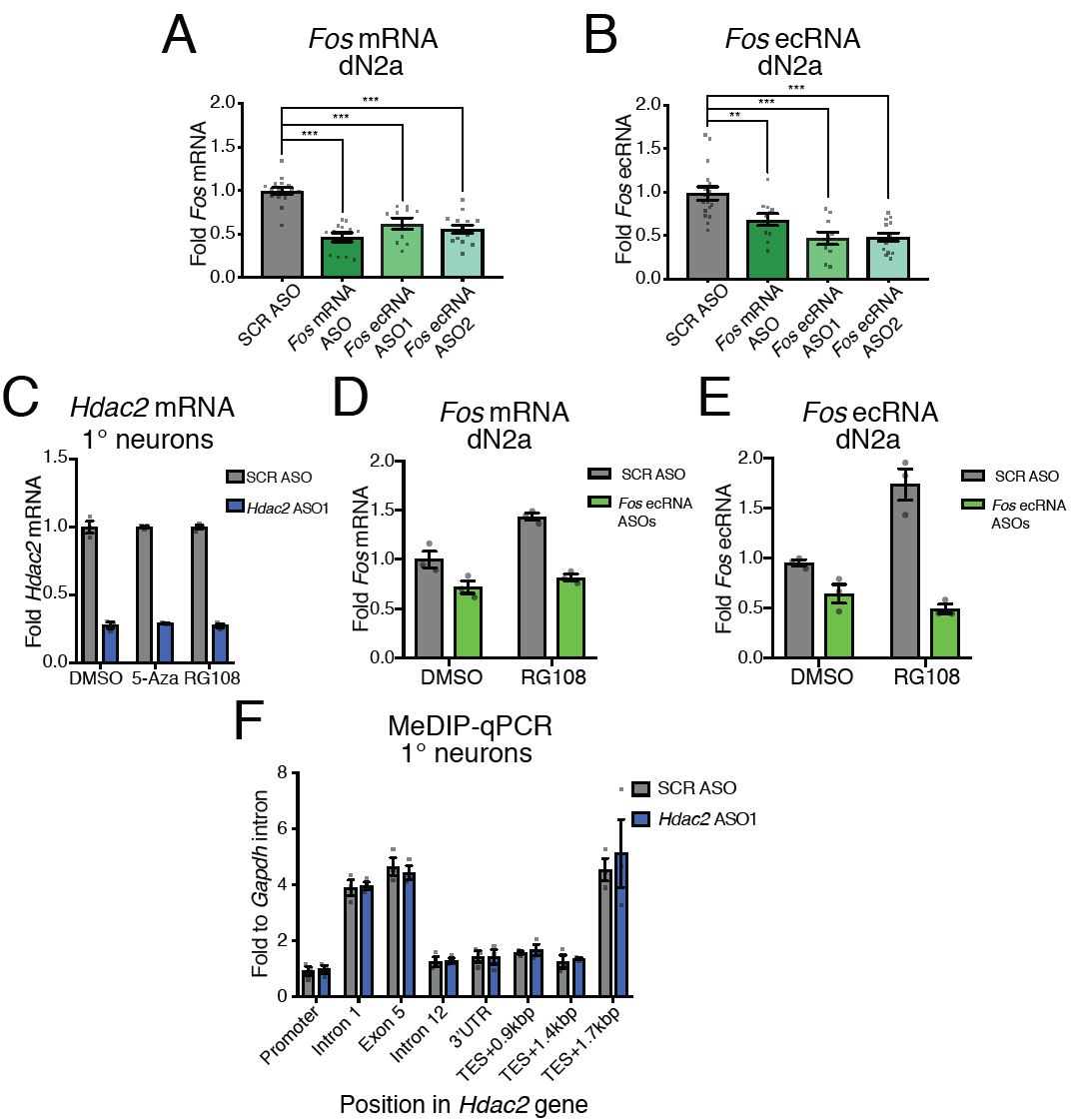

**Fig. S3. Effect of ASOs on ecRNA and DNA methylation.** (**A**) *Fos* mRNA expression after transfection of ASOs targeting *Fos* mRNA or *Fos* ecRNA. (**B**) *Fos* ecRNA expression after transfection of ASOs targeting *Fos* mRNA or *Fos* ecRNA. *N* = 10-17 wells of cell culture, one-way ANOVA with Dunnett’s multiple comparisons *post hoc* test. (**C**) RT-qPCR of *Hdac2* expression relative to *Hprt*. *N* = 3 wells of cell culture from technical replicates, fold change to SCR ASO of each treatment is plotted. Treated DNMT inhibitor for 1 day, then added ASO for 2 days, washed out 6 days before RNA was harvested. Doses were 5μM for 5-Aza and 200μM for RG108. No significant changes between *Hdac2* ASO1 treatments by One-way ANOVA. (**D**) *Fos* mRNA expression after transfection of Fos ecRNA ASOs (equimolar mix of both *Fos* ecRNA-targeting ASOs). 100μM RG108. (**E**) *Fos* ecRNA expression in same samples as panel D. (**F**) MeDIP-qPCR of primary cortical neurons treated with ASOs for 1-week. *N* = 3 biological replicates from independent preparations of primary neurons, averaged values from 3 technical replicates of the same experiment for each point on graph. Normalized to *Gapdh* intron primers and corresponding input sample. Two-way ANOVA was not below the level of significance. Bars represent ± SEM, grey dots show values for individual replicates. * = *p* < 0.05, ** = *p* < 0.01, *** = *p* < 0.001.

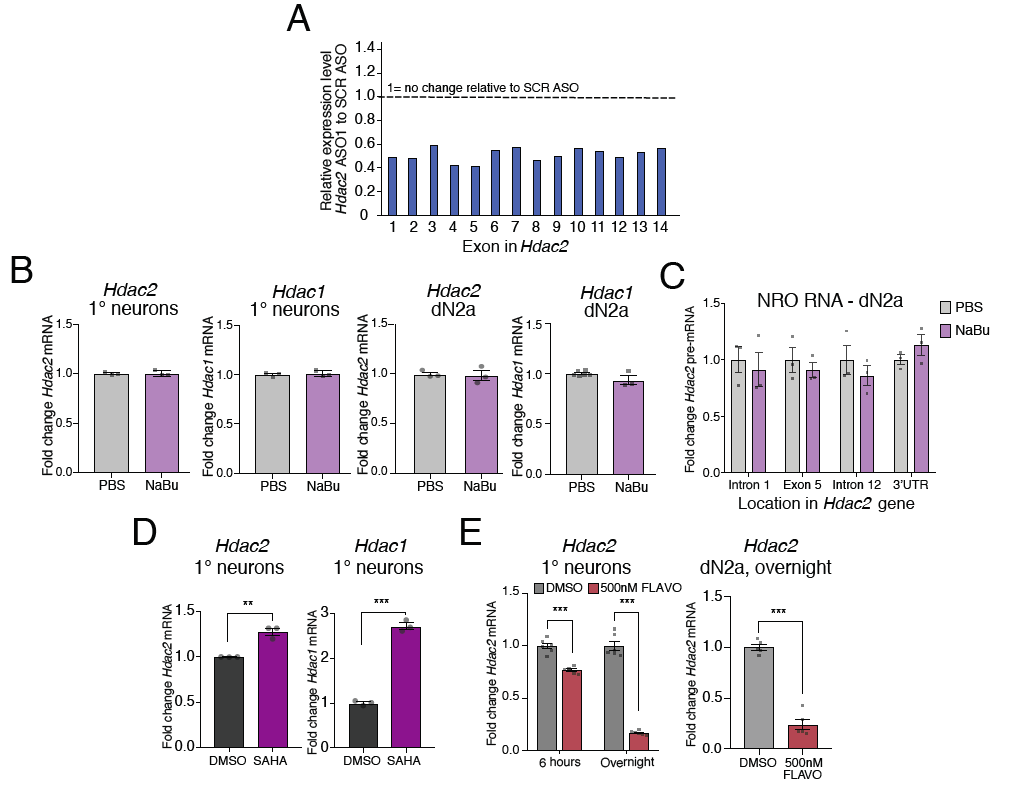

**Fig. S4. *R*educed transcriptional elongation at *Hdac2* after *Hdac2* ASO treatments in dN2a, HDAC inhibitors do not affect *Hdac2* transcription, and FLAVO represses *Hdac2* in dN2a.** (**A**) Fold change of *Hdac2* ASO1 animals relative to SCR ASO for each *Hdac2* exon was analyzed at 2 weeks after ICV injection in the hippocampus. (**B**) *Hdac2* and *Hdac1* mRNA expression in total RNA after 10μM NaBu treatment in primary cortical neurons and in dN2a. (**C**) NRO signal with indicated primers across *Hdac2* after 10μM NaBu treatment in dN2a. (**D**) *Hdac2* and *Hdac1* expression after 5μM SAHA treatment in primary neurons. (**E**) Primary neurons and dN2a were treated for 6 hours or overnight with 500nM FLAVO, and *Hdac2* expression was determined with RT-qPCR. RNA expression was normalized to *Hprt*. Student’s *t* tests were done for panels B, D, and E right (*N* = 3-5 replicates). Two-way ANOVA with Sidak’s multiple comparisons *post hoc* test was done in panel C (*N* = 3 technical replicates) and panel E left (*N* = 6 from 2 biological replicates done in triplicate). Error bars represent ± SEM, grey dots show values for individual replicates. * = *p* < 0.05, ** = *p* < 0.01, *** = *p* < 0.001.
